# Modelling potential range expansion of an underutilised food security crop in Sub-Saharan Africa

**DOI:** 10.1101/2021.09.15.460440

**Authors:** O. Koch, W.A. Mengesha, S. Pironon, T. Pagella, I. Ondo, I. Rosa, P. Wilkin, J.S. Borrell

## Abstract

Despite substantial growth in global agricultural production, food and nutritional insecurity is rising in Sub-Saharan Africa. Identification of underutilised indigenous crops with useful food security traits may provide part of the solution. Enset (*Ensete ventricosum*) is a perennial banana relative with cultivation restricted to southwestern Ethiopia, where high productivity and harvest flexibility enables it to provide a starch staple for ~20 million people. An extensive wild distribution suggests that a much larger region may be climatically suitable for cultivation. Here we use ensemble ecological niche modelling to predict the potential range for enset cultivation within southern and eastern Africa. We find contemporary bioclimatic suitability for a 12-fold range expansion, equating to 21.9% of crop land and 28.4% of the population in the region. Integration of crop wild relative diversity, which has broader climate tolerance, could enable a 19-fold expansion, particularly to dryer and warmer regions. Whilst climate change may cause a 37% – 52% reduction in potential range by 2070, large centres of suitability remain in the Ethiopian Highlands, Lake Victoria region and the Drakensberg Range. We combine our bioclimatic assessment with socioeconomic data to identify priority areas with high population density, seasonal food deficits and predominantly small-scale subsistence agriculture, where integrating enset may be particularly feasible and deliver climate resilience. When incorporating the genetic potential of wild populations, enset cultivation might prove feasible for an additional 87.2 - 111.5 million people, 27.7 – 33 million of which are in Ethiopia outside of enset’s current cultivation range. Finally, we consider explanations why enset cultivation has not expanded historically, and ethical implications of expanding previously underutilised species.

## 1. Introduction

Food and nutritional insecurity is a growing challenge in Sub-Saharan Africa (SSA) (Conceição *et al.* 2016; FAO and ECA 2018; Fraval *et al.* 2019), compounded by accelerating population growth, higher standards of living, degraded ecosystem services, climate change and volatile food markets (Poppy *et al.* 2014; Hall *et al.* 2017). Current efforts to address SSA food security through agricultural policies tend to emphasize increased productivity via inputs and technology (Conceição *et al.* 2016; Ittersum *et al.* 2016). However, a complementary strategy, which may be particularly pertinent under climate change, is the adaptation of agricultural systems through crop and cultivar choice (Rippke *et al.* 2016; J. S. Borrell *et al.* 2019; Mcmullin *et al.* 2019; Pironon *et al.* 2019; Rising and Devineni 2020). For example, recent evidence suggests that prioritising traits such as perenniality (Kreitzman *et al.* 2020), tolerance to drought or heat-induced stress (Heider *et al.* 2020) as well as crop diversity and asynchrony (Mcmullin *et al.* 2019; Renard and Tilman 2019; Egli *et al.* 2020) may help support smallholder resilience. Considering the antiquity and diversity of SSA agriculture, renewed investigation of orphan and underutilised indigenous crops may yield candidates with useful traits, where expanded cultivation could help meet food and nutritional security goals (Shelef *et al.* 2017; Tadele and Bartels 2019; Ulian *et al.* 2020).

In this study we investigate the indigenous Ethiopian food security crop enset (*Ensete ventricosum*, Musaceae), a close relative of the globally ubiquitous cultivated bananas (*Musa*). Enset provides the starch staple for 20 million Ethiopians (J S Borrell *et al.* 2019), and is one of 101 high potential crops identified by the African Orphan Crop Consortium (Dawson *et al.* 2018). Also known as the ‘false banana’, enset is a giant herbaceous perennial monocarp that accumulates standing biomass and can be harvested at any time prior to flowering and senescence (~7-12 years) (Lock 1993; National Research Council 2006). Upon harvesting, the entire pseudostem and corm is processed to extract starch, which is fermented and stored until required for consumption (Tamrat *et al.* 2020). Enset is non-irrigated and is among the highest yielding crops per hectare in the region, whilst vegetative propagation enables rapid multiplication of favourable genotypes (Borrell *et al.* 2020). By maintaining multiple age-classes, enset provides subsistence farmers the flexibility to harvest as required (e.g. depending on availability of other crops or resources), buffering seasonal, social and climate driven variability (J S Borrell *et al.* 2019). This suite of unusual food security traits has earned enset the moniker ‘the tree against hunger’ (Brandt *et al.* 1997). Nevertheless, despite its local agricultural dominance, utility and major cultural importance in the southwestern Ethiopian highlands, enset has a remarkably narrow cultivated distribution and is virtually unknown as a food plant outside of Ethiopia (J S Borrell *et al.* 2019).

Archaeological and historical evidence suggests that enset was domesticated in Ethiopia (Brandt *et al.* 1997; Hildebrand 2010) and that cultivation has remained restricted to the south-west (Negash 2020). There is limited evidence that cultivation was once more extensive in northern Ethiopia, as observed in Bruce (1790). By contrast, inedible wild enset populations are distributed across moist Afromontane Forest habitats in eastern and southern Africa (Borrell et al., 2019). This broad wild distribution provides an initial indication of the potential to expand domesticated enset cultivation beyond its current range. As a major African center of crop domestication, multiple Ethiopian crops including coffee (*Coffea arabica*) (Davis *et al.* 2018) and finger millet (*Eleusine coracana)* (Fuller 2014) have been successfully adopted beyond the species’ native range (Fuller and Lucas 2017). It is therefore surprising that as a regional staple and a close relative of the globally cultivated banana, enset has not been adopted outside of Ethiopia.

Climate change is predicted to seriously affect yields and distributions of major staple crops in Africa (Schlenker and Lobell 2010; Challinor *et al.* 2014; Pironon *et al.* 2019), which may catalyse renewed interest in adoption of alternative underutilised species (National Research Council 2006; Ulian *et al.* 2020; Mcmullin *et al.* 2021). Understanding barriers to adoption such as bioclimatic tolerance, prerequisite indigenous knowledge, access to material and opportunity costs is key to sustainably exploiting currently underutilised species (Bioversity International 2017; Jamnadass *et al.* 2020; Mcmullin *et al.* 2021). For example, a common attribute of cultivated species is reduced genetic diversity as a consequence of a domestication bottleneck (Gaut *et al.* 2018). Depending on the local bioclimatic conditions during the domestication process, genetic variants may have become fixed, limiting adaptive potential in parts of the species wild distribution (Warschefsky et al. 2014). Therefore, concurrently modelling the distribution of crop wild relative progenitor populations, from which genetic diversity could be introduced through breeding, may indicate further potential for crop expansion. Similarly, subsistence farmer agricultural practice and crop choice is strongly influenced by risk (Aryal *et al.* 2021), with the cost of crop adoption a trade-off against the risk of food insecurity or perceived climate vulnerability of their current crop portfolio (Akinyi et al. 2021).

In this study, we aim to identify both agrisystems and communities in which enset expansion may be appropriate, by characterising both the present and future bioclimatic distribution in which cultivation is viable, as well as the socioeconomic context in which adoption is feasible (Shikuku *et al.* 2017; Mcmullin *et al.* 2021). We apply an ensemble ecological niche modelling (ENM) framework to assess climatic suitability for expanded enset cultivation across eastern and southern Africa and integrate this with poverty, demography and food insecurity data to prioritise socioeconomic suitability. Relatively few studies have applied ecological niche modelling to evaluate expansion potential in cultivated species (e.g. baobab, Cuni and Patrick 2010; coffee, Moat et al. 2017), and fewer have attempted to integrate anthropogenic features (e.g. *Colocasia* in Hawaii, Kodis et al. 2018). Enset has significant potential because it can address acute food insecurity asset, even when grown in low numbers (National Research Council 2006) and the wild distribution is considerably larger than the current cultivated distribution (the inverse of many other domesticates, Diamond 2002).

We first generate ENMs for wild and domesticated enset across Ethiopia and assess niche shifts associated with domestication. We use these models to evaluate enset’s current and potential distribution, as well as explanations for the lack of historical expansion. Second, we evaluate the extent to which integrating wild enset genetic diversity could enable domesticated enset to adapt to a broader region of cultivation, both now and under future climate projections. Third, we analyse demographic and socioeconomic data to identify priority areas in which enset cultivation could contribute to food and nutritional insecurity needs with minimal barriers to adoption. Finally, we consider remaining political and cultural barriers to expansion of enset, and parallels of this approach with other underutilised species.

## 2. Methods

### Agricultural surveys and data processing

We collated 2515 georeferenced records of domesticated enset in the Ethiopian Highlands from 2017-20. Observations included systematic agricultural surveys oriented to transect elevational and climatic gradients from across the enset growing region, ensuring comprehensive coverage of variation in bioclimatic and geographical space. We also collated 163 wild *E. ventricosum* observations from across Ethiopia and East and Southern Africa using field surveys, online databases (GBIF.org 2018) and herbaria records (AAU, K).

We were cognisant that sampling bias in geographic space (e.g. due to accessibility) may translate to bias in environmental space and overfitting (Boria *et al.* 2014). To ensure a balanced representation of environmental conditions in our dataset, we filtered presence data in environmental space following Varela et al. (2014), using 19 bioclimatic predictors from CHELSA for the period 1979-2013 (Karger *et al.* 2017) and aggregated to 5 arc-minutes (~10 km^2^) resolution in ArcMap 10.7.1 (ESRI, USA). First, we plotted a Principal Component Analysis of scaled environmental variables for all observations using the R package ade4 (Dray and Dufour 2007), and removed occurrence points with duplicated environmental values on the first two principal components. This retained 414 domesticated and 99 wild enset occurrence points. We applied further bias correction to domesticated samples by calculating pairwise Euclidean distance and using Kmeans to identify 30 clusters that minimise within sums of squares. We randomly sampled three occurrence points from each, generating a sample evenly distributed across environmental space. We then partitioned 60 domesticated enset observations for model training and 30 for model evaluation from these clusters. Generating pseudo-absences (PAs) enables parameterization of modelling approaches that do not rely on true species absences. To account for uncertainty in PAs samples, we generated 10 datasets of 10,000 PAs randomly selected across the reference area, following the approach of Barbet-Massin et al. (2012).

### Variable selection and climate projections

We defined the study area as comprising Ethiopia (hereafter “reference area”) and the 17 countries of East and Southern Africa in which wild enset has been recorded (“transfer area”) (Figure 1). This modelling background extent was restricted to relevant agroecological zones to obtain informative evaluation statistics and model output (Lobo *et al.* 2012), using the “Agro-Ecological Zones for Africa South of the Sahara” dataset (HarvestChoice and International Food Policy Research Institute, 2015), omitting arid and sub-tropical regions. To reduce multicollinearity and model overfitting we restricted the number of environmental variables using the “select07” approach (Dormann *et al.* 2013). Where pairs of variables were correlated (Spearman’s rank correlation coefficient > 0.7), we retained the variable with the highest explanatory power by calculating the univariate importance of all 19 bioclimatic variables using the AIC of a quadratic GLM. Finally, ExDet (Mesgaran *et al.* 2014) was used to select a set of variables only moderately correlated in the reference area as well as the transfer area whilst also featuring low novelty in the transfer area. We therefore retained maximum temperature of the warmest month (Bio 5), precipitation of the driest quarter (Bio 17) and mean annual precipitation (Bio 12) as response variables (Table S1). We omitted edaphic variables as Ethiopian homegarden agriculture tends to alter local soil composition at fine scales (Wolka *et al.* 2021). Similarly, we consciously excluded other remotely sensed variables such as Normalized Difference Vegetation Index or Net Primary Production, as although these may improve present time models, it is unclear how they could be applied to future projections (Leitão *et al.* 2019).

**Figure 1.**
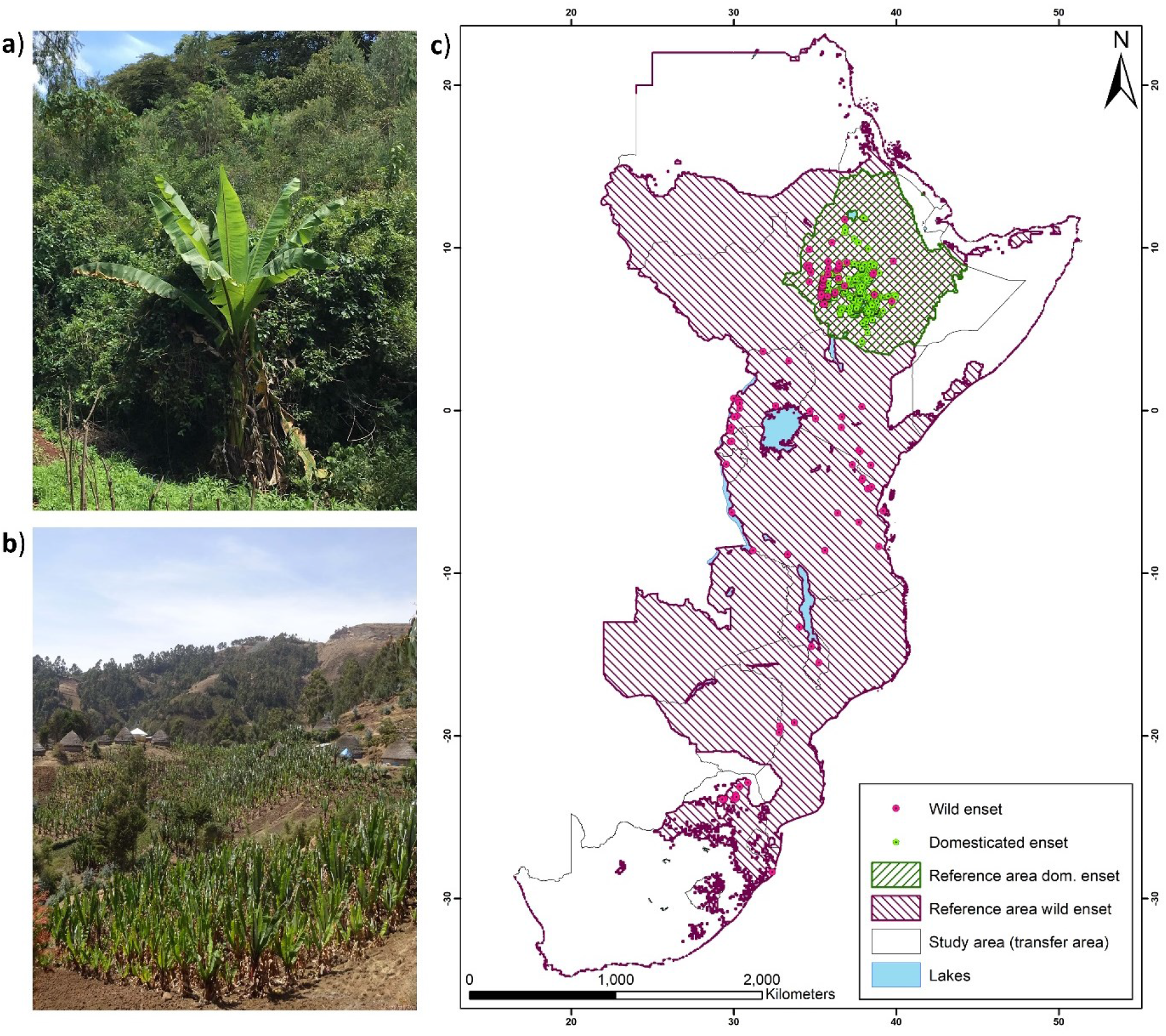
Extent of the study area in eastern and southern Africa and distribution data for wild and domesticated enset. a) Image of wild enset from a river valley near Bonga, Ethiopia. b) Cultivated enset from Basketo region, Ethiopia. c) Locations of 90 wild and 414 domesticated enset observations.

For modelling the effect of climate change on the area suitable for enset cultivation, projections of five global circulation models (GCMs) from the Coupled Model Intercomparison Project Phase 5 (CMIP5) (CESM1-BGC, CESM1-CAM5, CMCC-CM, MIROC5 and MPI-ESM-MR) were used for Representative Concentration Pathway Scenario (RCP) 4.5 and RCP8.5 in 2050 and 2070. GCM selection was based on dissimilarity following Sanderson et al. (2015), to reduce interdependence among the selected models.

### Ensemble modelling for wild and domesticated enset

We use an ensemble of six different modelling techniques: Generalized Linear Models (GLM), Generalized Additive Models (GAM), Generalized Boosting Models (GBM), Random Forest (RF), Multiple Adaptive Regression Splines (MARS) and Maximum Entropy (MAXENT), implemented in the R-package “biomod2” (Thuiller *et al.* 2016). Five modelling runs with different testing/training data splits were carried out on 10 different sets of pseudo-absences for each of the six modelling approaches, generating 300 models each for wild and domesticated enset. An ensemble was generated from all models with a true skill statistic (TSS) ≥0.6 and an area under the receiver operating characteristic curve ≥0.8 We combined models using the mean of probabilities weighted by each model’s TSS score. The maximized sum of sensitivity and specificity (equivalent to maximizing TSS) was used to generate binary presence-absence predictions (Barbet-Massin *et al.* 2012).

To compare domesticated and wild enset niches we used the niche similarity statistics Schoener’s D and Hellinger’s I (Warren *et al.* 2008), as well as testing niche expansion, unfilling and stability in the process of domestication using the ecospat package (Broennimann *et al.* 2021). To assess areas climatically suitable for enset cultivation outside of its current range, ensemble models were projected across the 18 countries of the study region under current and future climate conditions. Mean probabilities of occurrence were calculated across the five GCMs for each time period and RCP.

### Model evaluation

Ensemble model discrimination ability was evaluated using AUC and TSS on a 30% subsample of presences and pseudo-absences, across five independent runs. To evaluate calibration (ability to correctly predict conditional probability of presence), we plotted the continuous Boyce index (CBI) (Hirzel *et al.* 2006). The CBI is a threshold independent measure of model performance, which measures the distribution of observed presences across the projected suitability based on divergence from a random distribution. Good performance is indicated by an increasing number of presences found with increasing predicted probability of presence. In addition, we calculated the absolute validation index, a threshold dependent measurement of the proportion of presences correctly identified above the chosen binary probability of presence (Hirzel *et al.* 2006).

### Socioeconomic analysis of enset suitability

After identifying bioclimatic suitability for enset cultivation, we performed further prioritization based on five geographic, demographic, and socioeconomic criteria. First, to mitigate unsustainable agricultural expansion, we retained only crop land outside of IUCN category I/II protected areas, using the Spatial Production Allocation Model dataset for 2017 (International Food Policy Research Institute 2020) and World Database of Protected Areas (UNEP-WCMC and IUCN 2020). Second, enset is characterised by high harvest flexibility buffering seasonal food insecurity, therefore we integrated data from the Famine Early Warnings System Network (USAID, 2019) to identify areas of seasonal food access deficiencies from the period 2012-19. Average Integrated Food Security Phase Classification for January/February, June/July and October (Months where continuous timeseries were available) were obtained as a proxy for food access deficiencies per season. We retained areas with an average classification of two (“stressed”) or higher in any season.

Enset successfully supports very high rural population densities and has among the highest yield per hectare of regional crops (Borrell *et al.* 2020). Therefore third, we identified regions with high rural population density using WorldPop (2018), which may be indicative of current or future land shortages. The top 10^th^ (~10,000 people) and 5^th^ percentile (~18,000 people) of resampled cells were classified as densely populated and very densely populated, respectively. Fourth, enset is a low input, non-irrigated, predominantly subsistence cropping system. We used Spatially Disaggregated Crop Production Statistics Data in Africa South of the Sahara for 2017 (IFPRI, 2020) to identify areas with a high share of rain fed subsistence and low input cropping practices. The top 20^th^ (>~500 ha) and 10th percentile (>~1000 ha) of cells were classified as of priority and of high priority respectively. Finally, we used the same data to identify areas with low local diversity of staple crops. Crop diversity is an increasingly recognised strategy for mitigating food insecurity (Fraval *et al.* 2019; Koch *et al.* 2021). We calculated the Shannon Diversity Index for each cell based on the respective crops contributing to low input and subsistence production. The lowest 2/3 and 1/3 were classified as priority and high priority respectively. All indicators were scaled between 0-2 and the sum of indicator scores was used as an overall priority score of each cell. Priority maps were then overlaid with 2070 projections of binary suitability to account for climate change when prioritizing expansion areas. All analyses were performed in R software version 3.6.2 (R Core Team 2019).

## 3. Results

### Ensemble model performance

The domesticated enset ensemble model achieved a high discriminative ability with an overall TSS of 0.73 and AUC of 0.91. Individual model performance ranged from a TSS of 0.40-0.80 and AUC of 0.69-0.92. The performance of MARS and RF models was poorer compared to other model types (Figure 2). A CBI score of 0.97 confirmed the very high predictive ability of domesticated enset models in the reference area. Moreover, binary predictions of domesticated enset models in reference space detected 83.3% of evaluation data as true presences. The coefficient of variation between models was generally very low for moderate to highly suitable areas (Figure 2), but uncertainty was greater in areas with low suitability. Ensemble model performance for wild enset was similarly high with an overall TSS of 0.77, AUC of 0.94 and CBI value of 0.94 (Table S2).

**Figure 2.**
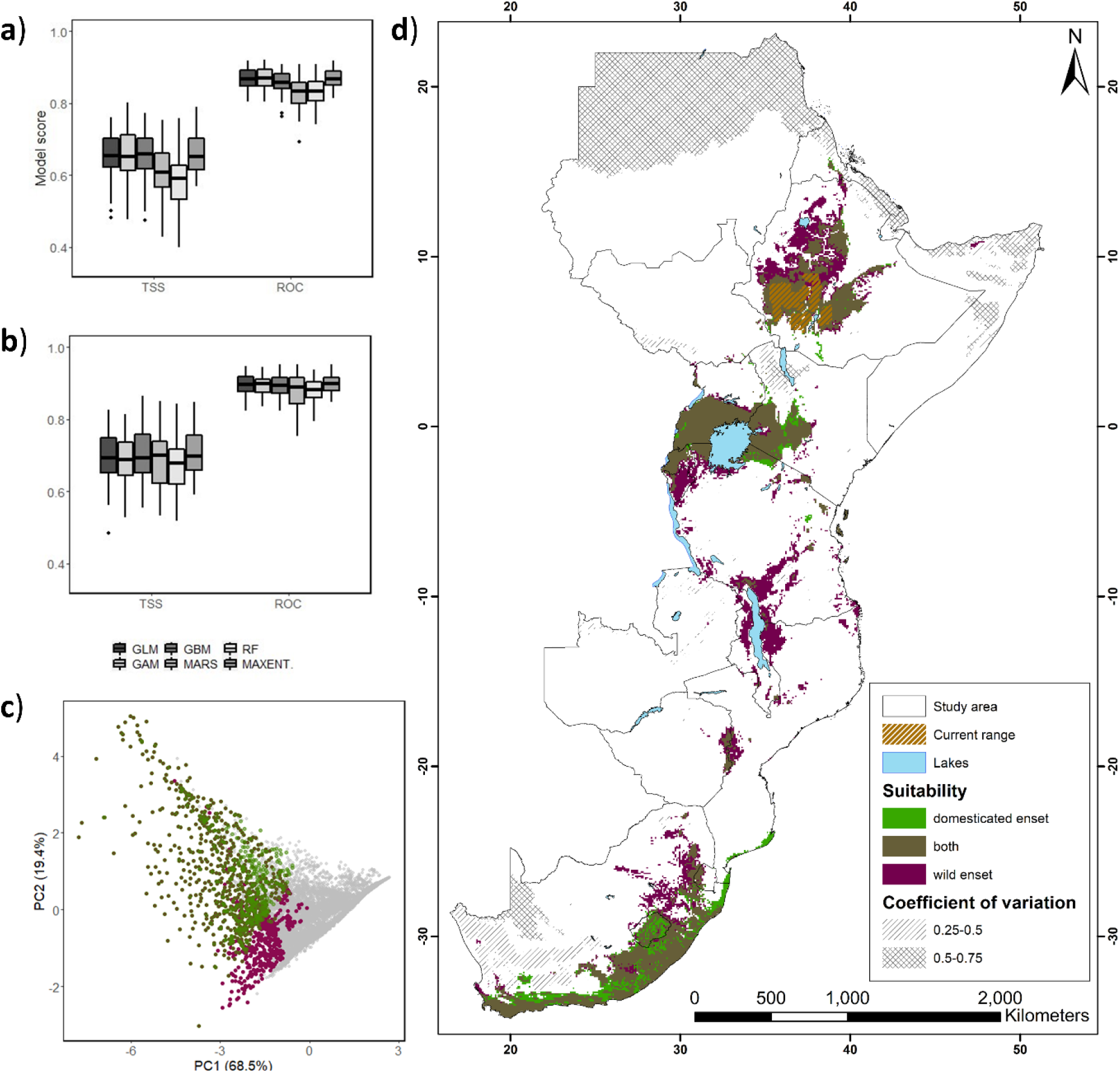
Potential suitability for domesticated and wild enset under current climate and expansion potential integrating the bioclimatic tolerance of wild enset populations. TSS and ROC scores across distribution modelling algorithms for a) domesticated and b) wild enset models. c) Comparison of the potential niche of enset in environmental space based on principal component analysis of bioclimatic variables. Grey denotes background points. d) binary representation of suitability, generated using the maximized TSS. The coefficient of variation shows the degree of uncertainty across model predictions.

### Comparison of enset’s wild and domesticated niches

Similarity statistics confirm substantial overlap between the wild and domesticated niche (D = 0.72, I = 0.89). However domesticated enset is confined to cooler maximum temperatures in the warmest month (Bio5) and driest quarter precipitation (Bio17) has consistently higher variable importance in ENMs (Figure 3). The expansion of domesticated enset’s niche is marginal (E=0.06%), while the unfilling of wild ensets niche is more pronounced (U= 6.5%). Together the maximum temperature of warmest month and the precipitation of the driest quarter have a combined relative importance of 98.5% for ensemble predictions, which offers a good representation of the climate suitability predicted for enset cultivation in the environmental space.

**Figure 3.**
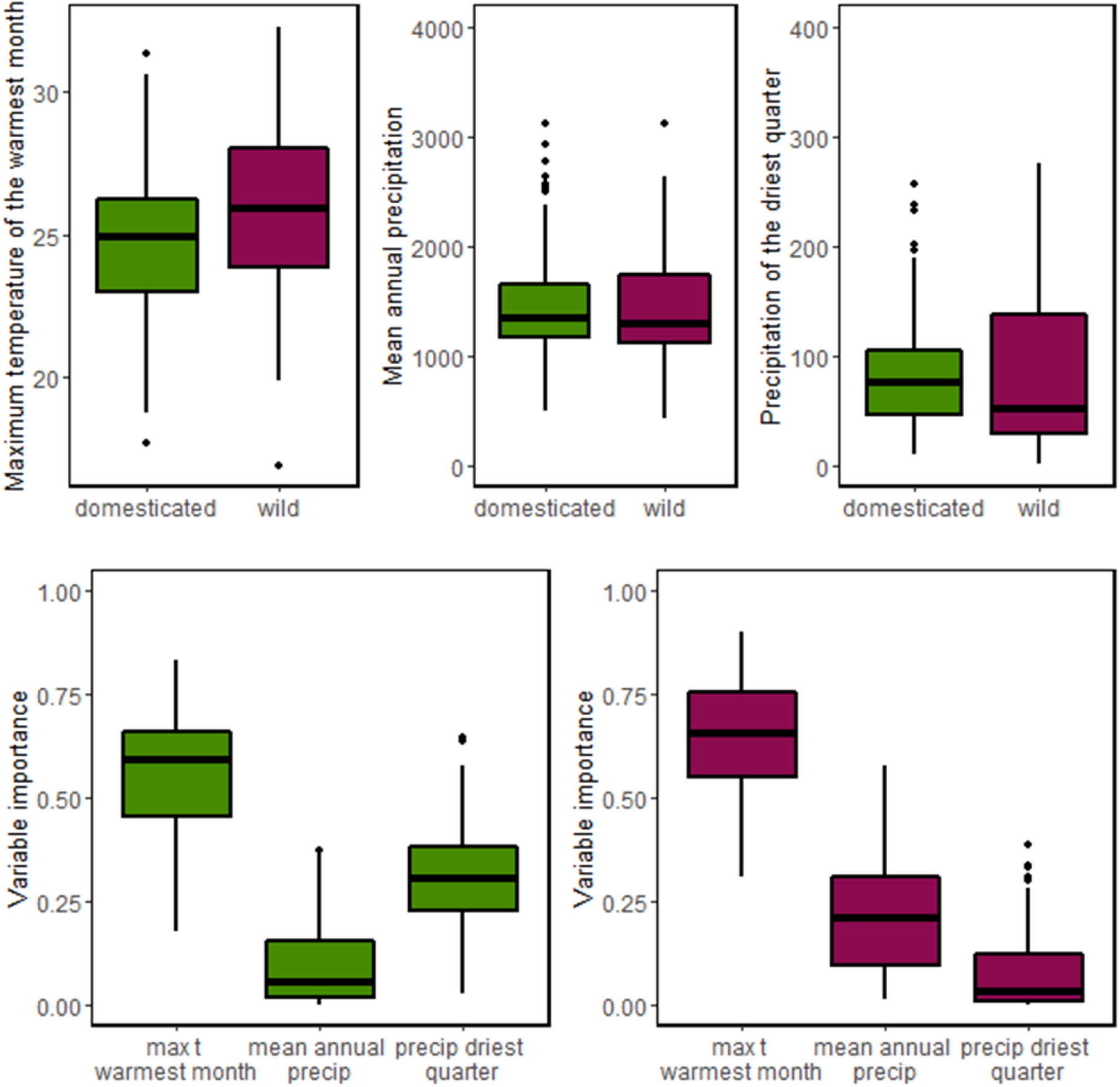
Comparison of the importance and range of bioclimatic variables for wild and domesticated enset. Bioclimatic values at enset presence points and model variable importance. While maximum temperature of the warmest month is significantly lower for domesticated enset (*t*(113) < 3.74, *p* < 0.001), differences in mean annual precipitation and driest quarter precipitation are non-significant.

### Potential for expansion of enset cultivation

Within Ethiopia, we estimate that approximately 251,300 km^2^ are within the range suitable for enset cultivation. Based on existing range maps from Borrell et al. (2019a), we estimate that ~28.2% of this extent is currently utilised. Across East and Southern Africa our ensemble model predicts ~906,000 km^2^ to be climatically suitable for enset cultivation, covering 21.9% (~208,000 km^2^) of croplands (Figure 2) and home to 28.4% of the total population of southern and eastern Africa. Country specific predictions are reported in Tables S3-5.

The wild enset ensemble model identified a substantially broader range of suitable areas (~1,270,000 km^2^), when compared to domesticated enset (906,000 km^2^). If we consider the potential for future breeding to integrate wild enset genetic diversity and the associated broader bioclimatic tolerance, ~1,375,000 km^2^ is suitable for enset, encompassing 37.8% (~361,000 km^2^) of the cropland in the study region. This represents a substantial further expansion than if domesticated enset is considered alone.

### Areas suitable for enset cultivation under projected future climate

Within its current Ethiopian range, enset cultivation is projected to contract by 10.8% under RCP4.5 and 20.9% under RCP8.5 by 2070. However, if we account for the potential to shift enset agriculture to all areas of suitability within Ethiopia, domestic enset’s range could be increased by 124.8% under RCP4.5 and 71.6% under RCP8.5. Across East and Southern Africa, both RCP4.5 and RCP8.5 scenarios show pronounced reduction in the range suitable for domesticated enset with respective declines of −31.3% and −47.4% by 2070 (Figure 4). Under projected future climates, the range of suitability for wild enset exceeds that of domesticated enset in most parts of the study area, thus the combined wild and domesticated enset model predictions reduce climate change induced range contractions of the potential enset cultivation area to −7% (RCP 4.5) and −29% (RCP 8.5). For 2050 scenarios see Figures S4 and S5.

**Figure 4.**
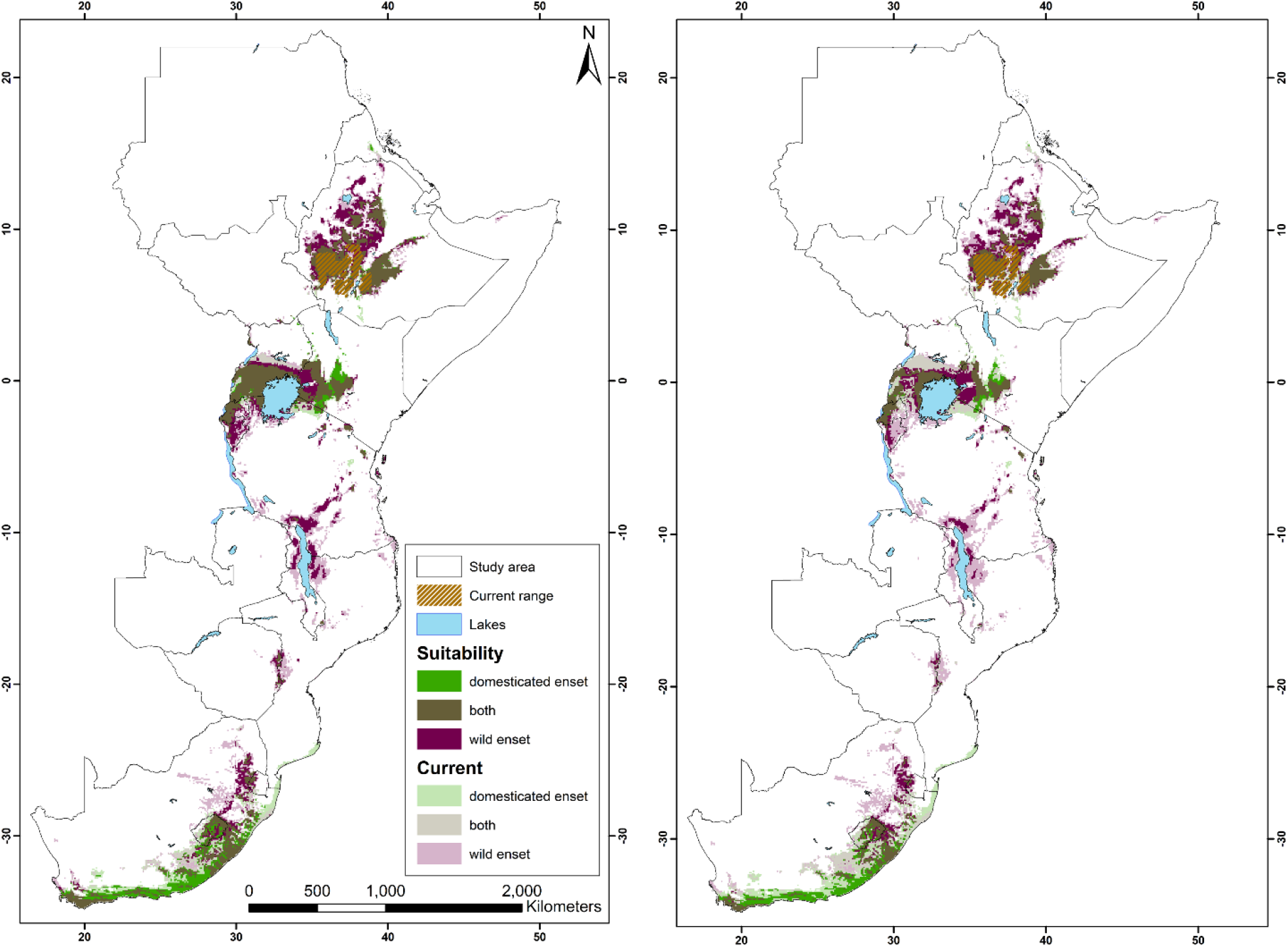
Suitability range change for wild and domesticated enset for 2070 under a) RCP4.5 and b) RCP8.5 scenarios. Projections for 2050 are provided in the supplementary materials figures S4 and S5.

### Socioeconomic prioritization of enset adoption

Using future bioclimatically suitable areas for 2070, we identified croplands characterised by low crop diversity, minimal agricultural inputs and high population density with frequent seasonal food security deficits. Based on this prioritization, we identify an additional current population of 12.8 – 19 million Ethiopians (depending on scenario) for which enset cultivation may be a beneficial, climate resilient asset to address food and nutritional insecurity outside its current cultivation area. More broadly across East and Southern Africa we identified 47 - 70.3 million people living in high priority areas, primarily localised to southern Uganda, eastern Kenya and Western Rwanda (see Figure 5). When incorporating the genetic potential of wild populations, enset cultivation might prove feasible for an additional 87.2 - 111.5 million people, 27.7 – 33 million of which are in Ethiopia outside of enset’s current cultivation range (see also Figures S6 and S7).

**Figure 5.**
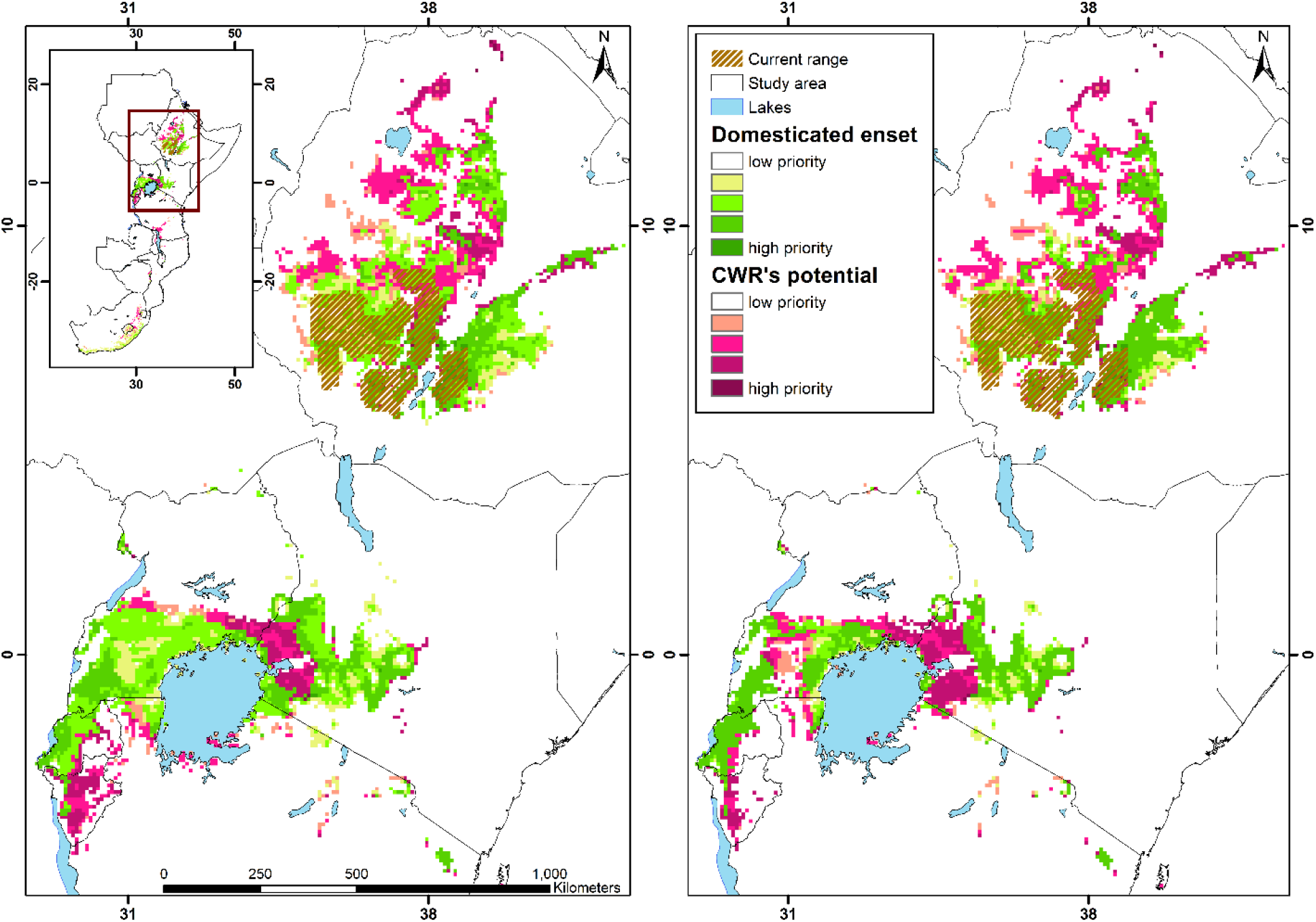
Priority areas for enset cultivation in east Africa based on demographic and socioeconomic metrics and enset suitability projections to 2070 under a) RCP 4.5 and b) RCP 8.5 scenarios. Both domesticated enset and the integration of crop wild relative diversity are considered. Region wide priority maps are provided in the supplementary materials figures S6 and S7.

## 4. Discussion

### Niche shifts associated with domestication

Comparison of wild and domesticated enset niches suggests that adaptation to on-farm cultivation shifted the realised niche towards cooler maximum temperatures and increased the importance of dry season precipitation. These changes are potentially indicative of the transition from growth in a shaded, moist Afromontane forest environment, where water availability is seasonally buffered, to cultivation in an open field environment. A requirement for cooler maximum temperatures may have resulted in the moderate eastward expansion of enset cultivation in Ethiopia to cropland at higher elevations. These bioclimatic limitations suggest that for domesticated enset, better emulating the conditions of its wild environment may contribute to broader bioclimatic tolerance, for example as a component of agroforestry systems, as practiced in some parts of Sidama zone (Abebe *et al.* 2010). Nevertheless, wild enset retains a broader overall niche space across all variables compared to domesticated enset, commensurate with presumed greater extant genetic diversity and absence of a domestication bottleneck. Surprisingly, the predicted suitability for both wild and domesticated enset across southwestern Ethiopia (Figure 2), suggests that the absence of contemporary wild populations across most of this range may have been due to extirpation, either through habitat conversion (e.g. loss of forest cover (Friis *et al.* 2010)), overexploitation (wild enset is inedible, but the leaves may be used (James S. Borrell *et al.* 2019)), lack of wild seed dispersers or genetic swamping from domesticated lineages.

Whilst domestication in many other plant species has been associated with expansion outside of their indigenous range (Diamond 2002), this has not occurred to any great extent in enset. Our models show that Ethiopia is isolated from other centres of enset suitability by at least 450 km, potentially resulting in a barrier to dispersal. Indeed, much of what is known about historic trade routes in Ethiopia suggests these were oriented towards Sudan and the Sahara as well as the Red Sea coast to Arabia where no suitable conditions exist for enset cultivation (Bent 1893; Pankhurst 1964). When combined with the additional barrier of indigenous knowledge associated with intensive enset cultivation, the processing requirements, and the fact that enset is propagated vegetatively which may travel less easily than seeds, this may partly explain why enset was not adopted elsewhere. Nevertheless, this would not have precluded concurrent domestication elsewhere in the wild range, as was likely the case for other crops such as rice in Asia (Civáň *et al.* 2015), though no evidence for use of enset in other cultures has been reported.

### Potential for expansion of enset cultivation

We find that despite a highly restricted current distribution, there is significant potential for climate-resilient enset expansion both within Ethiopia and across eastern and southern Africa (Figure 4). The closest areas with high suitability are in Amhara and Oromia regions of northern central Ethiopia. More widely, we also identify large areas of Kenya, Uganda and Rwanda which are characterised by a similar highland climate. Overall, ensemble model projections identified that 64.7% (~134,000 km^2^) of the cropland currently suitable for domesticated enset cultivation lies outside of Ethiopia under current climatic conditions.

The integration of genetic diversity and useful traits from wild progenitor populations and crop wild relatives is gaining increased attention as a pathway for climate change adaptation (Brozynska *et al.* 2016; Migicovsky and Myles 2017). Here we illustrate this value by showing that integrating crop wild relative diversity into enset breeding programmes may enable broader climate tolerance. Under current climate, this could enable expansion of enset cultivation by a further 144,000 km (15.1%) and under future scenarios up to ~73,000 km (7.6%) of the current cropland in the study area. Wild diversity may offer additional benefits beyond the potential for expansion. For example, higher fitness through improved tolerance of higher temperatures, even within existing bioclimatic limits, could translate into yield improvements (Zhao *et al.* 2017). Despite projected range declines for enset under high emission climate change scenarios, more than ~54,000 km^2^ (5.4%) of the current cropland are projected to remain suitable for enset cultivation outside of Ethiopia by 2070.

### Socioeconomic suitability for enset cultivation

Enset possesses a suite of traits that buffer acute food insecurity. We combined our climate data with population, food insecurity and agricultural inputs data to identify regions and communities with a similar socioeconomic context to those where enset is currently successfully utilised. This approach revealed priority areas in Ethiopia, as well as Kenya, Uganda and Rwanda with high future climate suitability, high rural population densities, frequent seasonal food deficits, low agricultural inputs and low current crop diversity. While achieving zero hunger is a major sustainable development goal (SDG2), agricultural expansion risks undermining related global efforts to reduce biodiversity loss (Molotoks *et al.* 2017). Highly flexible and productive species such as enset provide one pathway for improving local food security while minimising cropland expansion and resulting biodiversity loss, particularly because environmental degradation may be highest during periods of acute vulnerability, food insecurity and associated poverty (Asefa 2003).

### Remaining barriers to enset expansion

Even if barriers are low, the uptake of novel cultivation practices represents a risk, particularly for subsistence farmers (Meijer *et al.* 2015). Therefore, the bioclimatic and socioeconomic matching performed here, offers no guarantee that farmers will perceive and experience benefits. Previous approaches to disseminating agricultural innovations have focused on demonstration farms, farmer-to-farmer learning and *in situ* inclusive development of new approaches (Meijer *et al.* 2015), as part of extensive research on smallholder climate adaptation (Bryan *et al.* 2009; Conway and Schipper 2011; Shikuku *et al.* 2017). Successful examples include cassava (*Manihot esculenta*), which has expanded within Zambia to help mitigate drought vulnerability and associated food shortages (Barratt *et al.* 2006). However, we identify two remaining barriers that will require social and political approaches to overcome. First, Ethiopia currently restricts the international transfer of plant material to protect indigenous bioresources from inequitable exploitation (Tesgera 2019). Thus expansion outside of Ethiopia would depend on bilateral Access and Benefit Sharing Agreements, the implementation of which is highly variable internationally (Robinson *et al.* 2020). Second, the contemporary distribution of enset cultivation is currently closely associated with cultural groups who hold the required knowledge (Olango *et al.* 2014). This highlights that both knowledge as well as plant material would need to be fairly and equitably shared for successful transfer of enset cultivation (Swiderska 2006).

## Conclusions

Expanding the range of cultivation of currently underutilised crops has significant potential to support the diversification and resilience of global agrisystems under climate change. Unifying interdisciplinary approaches involving both bioclimatic and socioeconomic suitability may help prioritise communities for agricultural development interventions, making successful adoption more likely. Whilst this represents a challenge to existing agrisystem and food networks, it is also an opportunity to adopt and improved suite of climate-resilient crops with multiple food security co-benefits.

## Funding and Acknowledgements

This work was supported by the GCRF Foundation Awards for Global Agricultural and Food Systems Research, entitled, ‘Modelling and genomics resources to enhance exploitation of the sustainable and diverse Ethiopian starch crop enset and support livelihoods’ [Grant No. BB/P02307X/1] and the GCRF I-FLIP grant ‘Enhancing enset agriculture with mobile agri-data, knowledge interchange and climate adapted genotypes to support the Enset Center of Excellence’ [BB/S018980/1]. JB was additionally supported by a Future Leader Fellowship at the Royal Botanic Gardens, Kew.

## Author contributions

OK and JB designed the study. JB and WA collected field observation. OK performed the analyses, with SP, IO and JB providing technical input. JB and PW secured funding. JB provided supervision. OK prepared the first draft and all authors contributed to the interpretation of the results and preparation of the final manuscript.

## Data Availability

Any data that support the findings of this study as well as the generated spatial data are available on Figshare (https://doi.org/10.6084/m9.figshare.16455648) under the CC BY 4.0 licence.

