## Supplementary material for "Modelling potential range expansion of an underutilised food security crop in Sub-Saharan Africa": Suplementary_Methods_Tables_Figures

Supplementary Figures ........ 1

### Supplementary Methods

#### *Priority area identification - Staple crop diversity*

We calculated the Shannon Diversity Index (H) for each cell based on the respective crops contributing to low input and subsistence production. H was calculated as follows:

$H = -\Sigma p\mathrm{i} * \ln(p\mathrm{i})$

Where $p$i describes the proportion the production of crop i contributes to the total crop production of a cell. The “overlay” function of the raster package (Hijmans et al., 2021) was used for the calculation of the stacked crop production raster layers. The production (measured in metric tonnes) in rainfed subsistence and low input cultivation of the crop groups “cereal”, “roots”, “pulses”, “millet” in the SPAM 2017 data set (HarvestChoice and International Food Policy Research Institute 2020) as well as plantain of the group “banana + plantains” and “rest of crops” were considered for staple crops (21 in total).

Supplementary Tables

**Table S1.** Selection process of bioclimatic variables with the list of retained variables after each step.

| 1. Initial set | 2. "select07" | 3. Transferability |
|---|---|---|
| Annual Mean Temperature (Bio 1) | | |
| Mean Diurnal Range (Bio 2) | | |
| Isothermality (Bio 3) | | |
| Temperature Seasonality (Bio 4) | X | |
| Max Temperature of Warmest Month (Bio 5) | X | **X** |
| Min Temperature of Coldest Month (Bio 6) | | |
| Temperature Annual Range (Bio 7) | | |
| Mean Temperature of Wettest Quarter (Bio 8) | | |
| Mean Temperature of Driest Quarter (Bio 9) | | |
| Mean Temperature of Warmest Quarter (Bio 10) | | |
| Mean Temperature of Coldest Quarter (Bio 11) | | |
| Annual Precipitation (Bio 12) | X | **X** |
| Precipitation of Wettest Month (Bio 13) | | |
| Precipitation of Driest Month (Bio 14) | | |
| Precipitation Seasonality (Bio 15) | | |
| Precipitation of Wettest Quarter (Bio 16) | | |
| Precipitation of Driest Quarter (Bio 17) | X | **X** |
| Precipitation of Warmest Quarter (Bio 18) | | |
| Precipitation of Coldest Quarter (Bio 19) | X | |

**Table S2** Model evaluation and validation scores for ensemble models on domesticated and wild enset.

| | Model evaluation | | | Model validation | |
|---|---|---|---|---|---|
| **Modelled species** | **TSS** | **AUC** | **Sensitivity** | **CBI** | **AVI** |
| domesticated *E. ventricosum* | 0.947 | 0.995 | 97.5 | 0.937 | - |
| wild *E. ventricosum* | 0.957 | 0.997 | 97.778 | 0.893 | 0.66 |

**Table S3. Range suitable for enset cultivation and total area of cropland across the study area for current climate**. Countries without suitable ranges throughout current and future climates are not listed.

| Current | Domesticated enset | | | | Domesticated and wild enset | | | |
|---|---|---|---|---|---|---|---|---|
| | potential range | | cropland | | potential range | | cropland | |
| Country | area | share | area | share | area | share | area | share |
| Zimbabwe | 8000 | 1.7% | 802 | 2.5% | 14300 | 3.0% | 1580 | 4.9% |
| Zambia | 0 | 0.0% | 0 | 0.0% | 3600 | 0.4% | 448 | 1.2% |
| South Africa | 313800 | 19.3% | 10501 | 13.6% | 403300 | 24.8% | 19580 | 25.4% |
| Uganda | 110200 | 45.7% | 33802 | 55.3% | 116100 | 48.1% | 36071 | 59.0% |
| Tanzania | 54300 | 5.2% | 19531 | 9.3% | 158600 | 15.2% | 52308 | 25.0% |
| Swaziland | 6800 | 29.4% | 749 | 29.7% | 8500 | 36.8% | 969 | 38.5% |
| South Sudan | 400 | 0.1% | 8 | 0.0% | 1300 | 0.2% | 316 | 1.0% |
| Rwanda | 20200 | 70.1% | 14510 | 69.0% | 25200 | 87.5% | 20950 | 99.6% |
| Malawi | 1800 | 1.6% | 1380 | 1.7% | 23600 | 20.3% | 12903 | 15.5% |
| Mozambique | 12200 | 1.3% | 1844 | 2.2% | 50000 | 5.2% | 9418 | 11.2% |
| Lesotho | 29700 | 72.4% | 1773 | 65.1% | 31200 | 76.1% | 1952 | 71.7% |
| Kenya | 90400 | 13.4% | 45422 | 53.4% | 98200 | 14.6% | 51598 | 60.7% |
| Ethiopia | 251300 | 18.9% | 74186 | 40.6% | 393600 | 29.5% | 130227 | 71.3% |
| Eritrea | 3400 | 2.4% | 1609 | 16.6% | 4300 | 3.0% | 1967 | 20.3% |
| Burundi | 3400 | 11.5% | 2582 | 11.1% | 23900 | 80.7% | 20365 | 87.6% |
| Study Area | 905,900 | 8.7% | 208,700 | 21.9% | 1,355,700 | 13.0% | 360,654 | 37.9% |

**Table S4. Range suitable for enset cultivation and total area of cropland across the study area for climate in 2050**. Countries without suitable ranges throughout current and future climates are not listed, and we assume cropland remains static.

| 2050 | RCP 4.5 | | | | | | | | RCP 8.5 | | | | | | | |
|---|---|---|---|---|---|---|---|---|---|---|---|---|---|---|---|---|
| | Domesticated enset | | | | Domesticated and wild enset | | | | Domesticated enset | | | | Domesticated and wild enset | | | |
| | potential range | | cropland | | potential range | | cropland | | potential range | | cropland | | potential range | | cropland | |
| **Country** | **area** | **share** | **area** | **share** | **area** | **share** | **area** | **share** | **area** | **share** | **area** | **share** | **area** | **share** | **area** | **share** |
| Zimbabwe | 3300 | 0.7% | 161 | 0.5% | 7100 | 1.5% | 596 | 1.8% | 2000 | 0.4% | 87 | 0.3% | 5600 | 1.2% | 366 | 1.1% |
| Zambia | 0 | 0.0% | 0 | 0.0% | 0 | 0.0% | 0 | 0.0% | 0 | 0.0% | 0 | 0.0% | 0 | 0.0% | 0 | 0.0% |
| South Africa | 206500 | 12.7% | 6549 | 8.5% | 246400 | 15.1% | 8415 | 10.9% | 167400 | 10.3% | 5672 | 7.4% | 206800 | 12.7% | 7300 | 9.5% |
| Uganda | 86900 | 36.0% | 25392 | 41.6% | 99800 | 41.4% | 31007 | 50.7% | 79300 | 32.9% | 23882 | 39.1% | 91500 | 37.9% | 28708 | 47.0% |
| Tanzania | 28000 | 2.7% | 12027 | 5.7% | 77200 | 7.4% | 28369 | 13.6% | 19100 | 1.8% | 9292 | 4.4% | 61900 | 5.9% | 23740 | 11.3% |
| Swaziland | 2700 | 11.7% | 303 | 12.0% | 5200 | 22.5% | 545 | 21.7% | 2100 | 9.1% | 210 | 8.3% | 5000 | 21.6% | 498 | 19.8% |
| South Sudan | 200 | 0.0% | 0 | 0.0% | 400 | 0.1% | 8 | 0.0% | 300 | 0.0% | 9 | 0.0% | 400 | 0.1% | 10 | 0.0% |
| Rwanda | 20700 | 71.9% | 15001 | 71.4% | 22800 | 79.2% | 16813 | 80.0% | 14900 | 51.7% | 11122 | 52.9% | 18600 | 64.6% | 14236 | 67.7% |
| Malawi | 500 | 0.4% | 131 | 0.2% | 6200 | 5.3% | 1550 | 1.9% | 500 | 0.4% | 131 | 0.2% | 5300 | 4.6% | 1262 | 1.5% |
| Mozambique | 300 | 0.0% | 13 | 0.0% | 7800 | 0.8% | 500 | 0.6% | 300 | 0.0% | 13 | 0.0% | 6200 | 0.6% | 371 | 0.4% |
| Lesotho | 23000 | 56.1% | 774 | 28.4% | 25400 | 62.0% | 1009 | 37.1% | 17500 | 42.7% | 326 | 12.0% | 22800 | 55.6% | 736 | 27.0% |
| Kenya | 69300 | 10.3% | 33439 | 39.3% | 88300 | 13.1% | 46897 | 55.1% | 65800 | 9.8% | 31608 | 37.2% | 84400 | 12.5% | 45441 | 53.4% |
| Ethiopia | 197500 | 14.8% | 58216 | 31.9% | 312300 | 23.4% | 101868 | 55.7% | 203400 | 15.3% | 60897 | 33.3% | 300400 | 22.6% | 97829 | 53.5% |
| Eritrea | 400 | 0.3% | 305 | 3.2% | 600 | 0.4% | 408 | 4.2% | 600 | 0.4% | 470 | 4.9% | 600 | 0.4% | 470 | 4.9% |
| Burundi | 3700 | 12.5% | 2643 | 11.4% | 17800 | 60.1% | 15368 | 66.1% | 1800 | 6.1% | 997 | 4.3% | 13500 | 45.6% | 12047 | 51.8% |
| Study Area | 643,000 | 6.2% | 154,954 | 16.3% | 917,300 | 8.8% | 253,352 | 26.6% | 575,000 | 5.5% | 144,716 | 15.2% | 823,000 | 7.9% | 233,014 | 24.5% |

**Table S5. Range suitable for enset cultivation and total area of cropland across the study area for climate in 2070**. Countries without suitable ranges throughout current and future climates are not listed, and we assume cropland remains static.

| **2070** | **RCP 4.5** | | | | | | | | **RCP 8.5** | | | | | | | |
|---|---|---|---|---|---|---|---|---|---|---|---|---|---|---|---|---|
| | **Domesticated enset** | | | | **Domesticated and wild enset** | | | | **Domesticated enset** | | | | **Domesticated and wild enset** | | | |
| | **potential range** | | **cropland** | | **potential range** | | **cropland** | | **potential range** | | **cropland** | | **potential range** | | **cropland** | |
| **Country** | **area** | **share** | **area** | **share** | **area** | **share** | **area** | **share** | **area** | **share** | **area** | **share** | **area** | **share** | **area** | **share** |
| Zimbabwe | 3000 | 0.6% | 146 | 0.4% | 7300 | 1.5% | 622 | 1.9% | 300 | 0.1% | 9 | 0.0% | 3600 | 0.8% | 167 | 0.5% |
| Zambia | 0 | 0.0% | 0 | 0.0% | 0 | 0.0% | 0 | 0.0% | 0 | 0.0% | 0 | 0.0% | 0 | 0.0% | 0 | 0.0% |
| South Africa | 175800 | 10.8% | 5893 | 7.6% | 214200 | 13.2% | 7641 | 9.9% | 129100 | 7.9% | 4383 | 5.7% | 160700 | 9.9% | 5387 | 7.0% |
| Uganda | 78400 | 32.5% | 23647 | 38.7% | 91500 | 37.9% | 29035 | 47.5% | 43100 | 17.9% | 13880 | 22.7% | 66200 | 27.4% | 21992 | 36.0% |
| Tanzania | 21400 | 2.0% | 9809 | 4.7% | 68000 | 6.5% | 25463 | 12.2% | 9100 | 0.9% | 4649 | 2.2% | 34700 | 3.3% | 13914 | 6.7% |
| Swaziland | 2400 | 10.4% | 260 | 10.3% | 5000 | 21.6% | 498 | 19.8% | 1100 | 4.8% | 103 | 4.1% | 3700 | 16.0% | 380 | 15.1% |
| South Sudan | 200 | 0.0% | 0 | 0.0% | 300 | 0.0% | 1 | 0.0% | 100 | 0.0% | 0 | 0.0% | 200 | 0.0% | 1 | 0.0% |
| Rwanda | 14600 | 50.7% | 11118 | 52.9% | 19600 | 68.1% | 15086 | 71.8% | 12600 | 43.8% | 9445 | 44.9% | 14900 | 51.7% | 11739 | 55.8% |
| Malawi | 500 | 0.4% | 131 | 0.2% | 5900 | 5.1% | 1444 | 1.7% | 300 | 0.3% | 52 | 0.1% | 1900 | 1.6% | 230 | 0.3% |
| Mozambique | 300 | 0.0% | 13 | 0.0% | 7600 | 0.8% | 474 | 0.6% | 100 | 0.0% | 8 | 0.0% | 1500 | 0.2% | 61 | 0.1% |
| Lesotho | 18400 | 44.9% | 357 | 13.1% | 23000 | 56.1% | 737 | 27.1% | 15600 | 38.0% | 184 | 6.8% | 20200 | 49.3% | 536 | 19.7% |
| Kenya | 65400 | 9.7% | 30653 | 36.0% | 86000 | 12.8% | 45295 | 53.3% | 46200 | 6.9% | 20491 | 24.1% | 73300 | 10.9% | 38613 | 45.4% |
| Ethiopia | 197800 | 14.8% | 58891 | 32.2% | 297400 | 22.3% | 95046 | 52.0% | 151000 | 11.3% | 43719 | 23.9% | 250200 | 18.8% | 76530 | 41.9% |
| Eritrea | 700 | 0.5% | 565 | 5.8% | 700 | 0.5% | 565 | 5.8% | 100 | 0.1% | 91 | 0.9% | 100 | 0.1% | 91 | 0.9% |
| Burundi | 1900 | 6.4% | 1040 | 4.5% | 16100 | 54.4% | 14246 | 61.3% | 1000 | 3.4% | 531 | 2.3% | 7700 | 26.0% | 6164 | 26.5% |
| Study Area | 580,800 | 5.6% | 142,523 | 15.0% | 842,600 | 8.1% | 236,153 | 24.8% | 409,700 | 3.9% | 97,546 | 10.3% | 638,900 | 6.1% | 175,805 | 18.5% |

Supplementary Figures

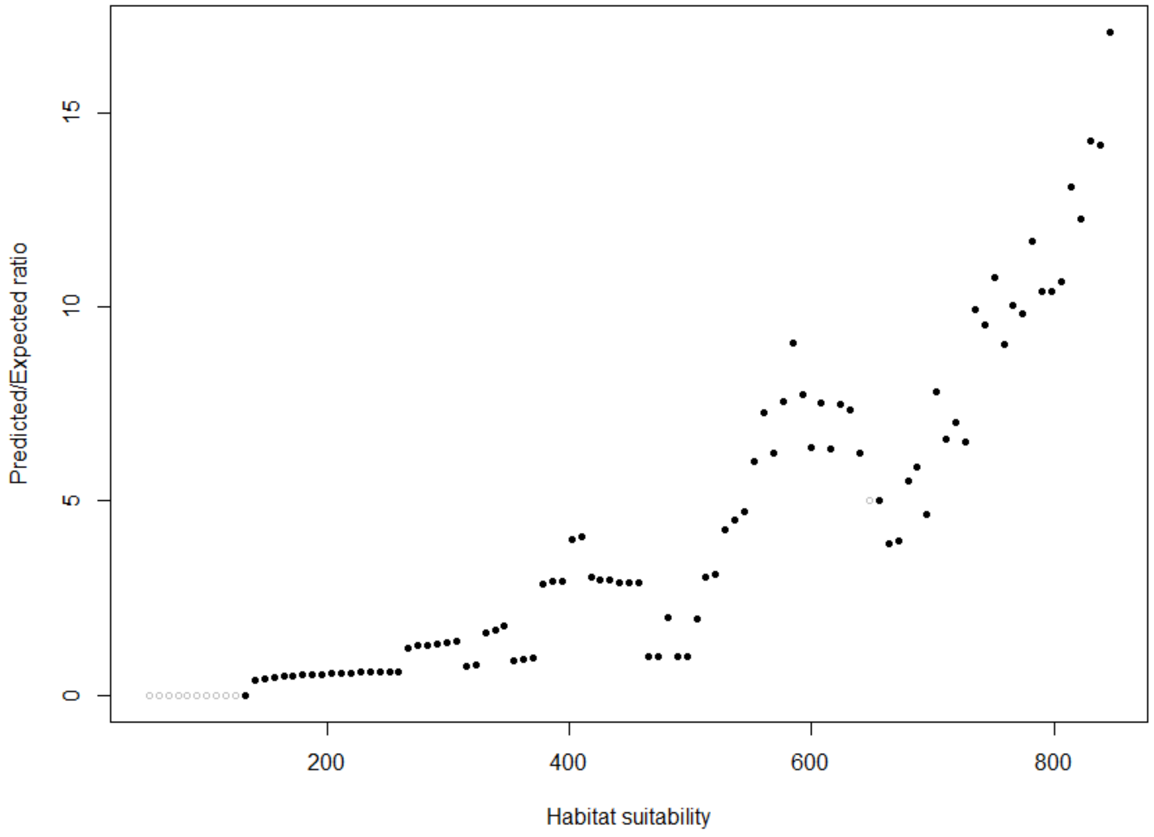

**Figure S1.** Boyce index plot of domesticated enset ensemble model predictions and validation points in the reference area.

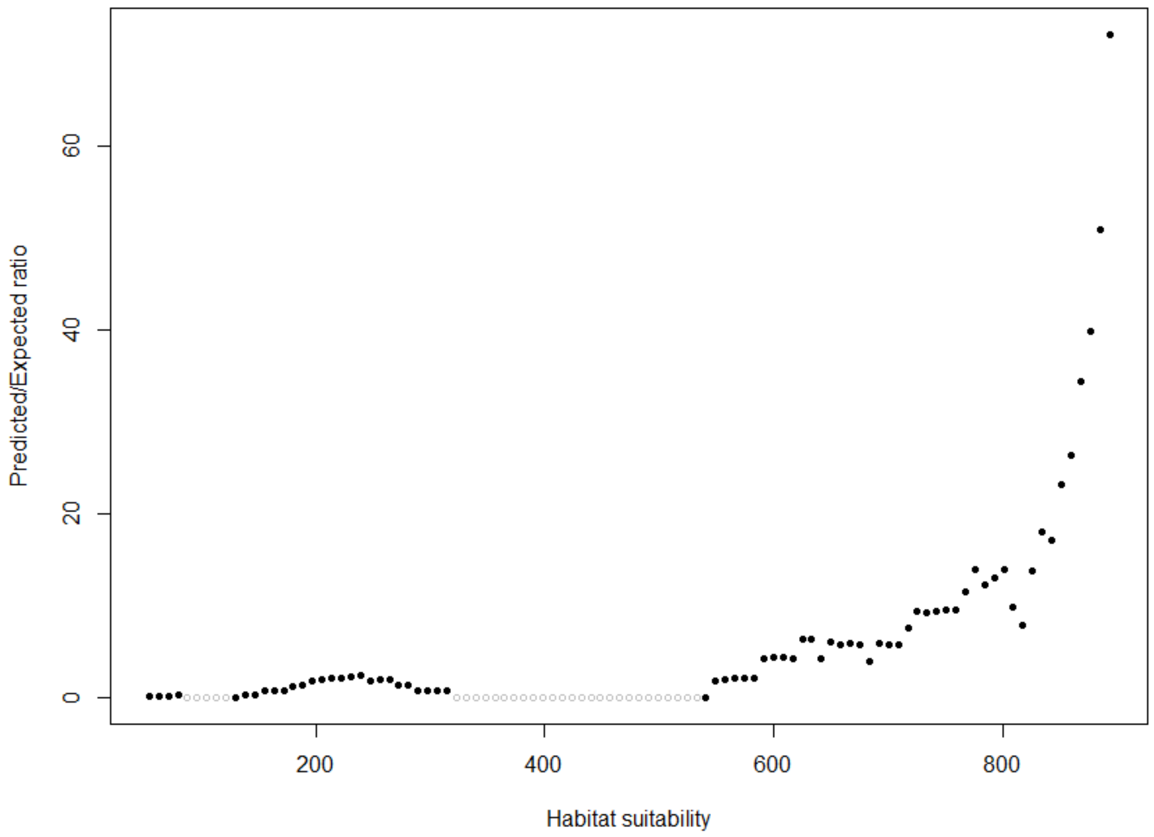

**Figure S2.** Boyce index plot of wild enset ensemble model predictions and validation points in the respective reference area.

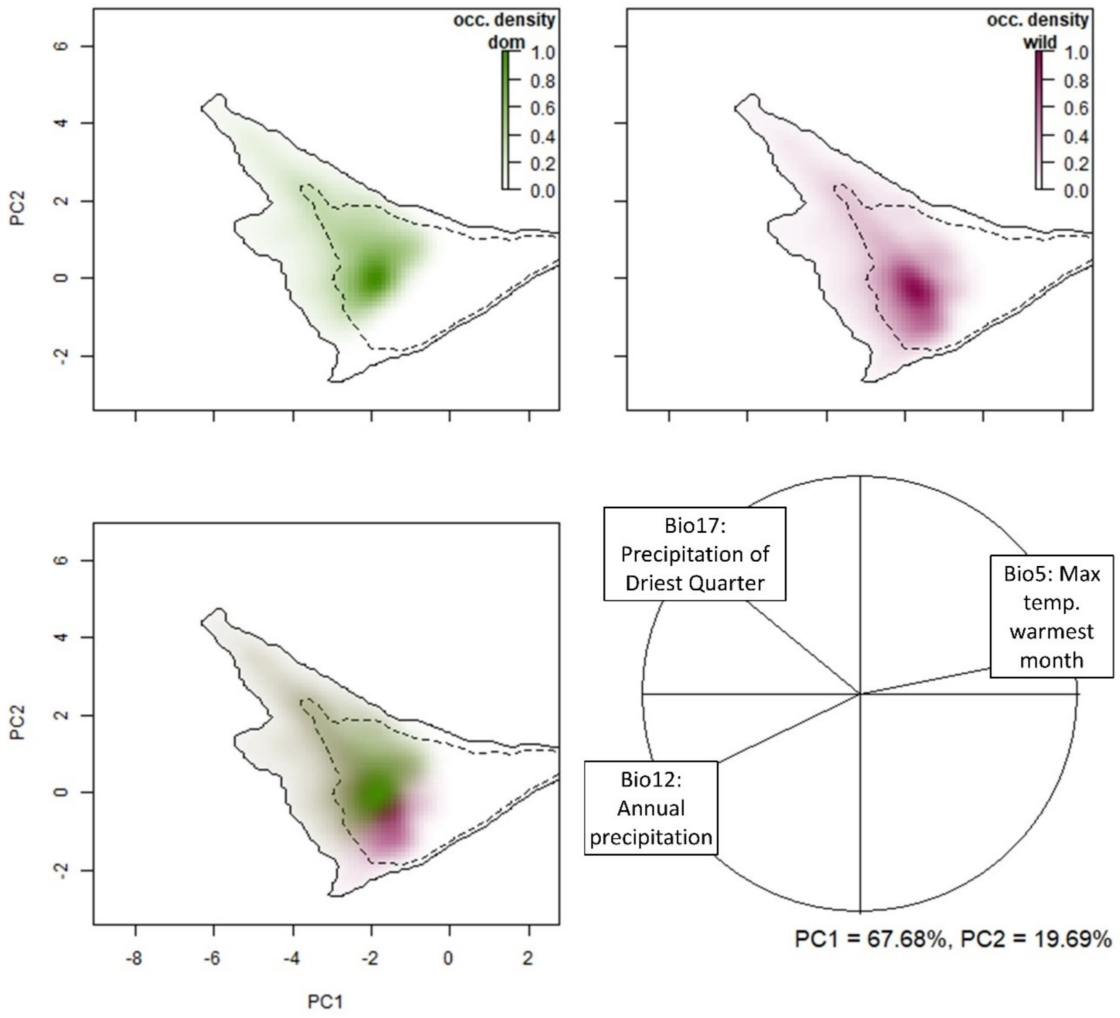

**Figure S3. Comparison of the potential niche of enset in the study area based on principal component analysis of bioclimatic variables**. Colour saturation represents the density of the species occurrence per cell (green, domesticated enset; purple, wild enset). Solid line represents 99.5% and dashed line represents 50% of the available background. Variables comprise Bio5 = Max Temperature of Warmest Month; Bio12 = Annual Precipitation and Bio17 = Precipitation of Driest Quarter.

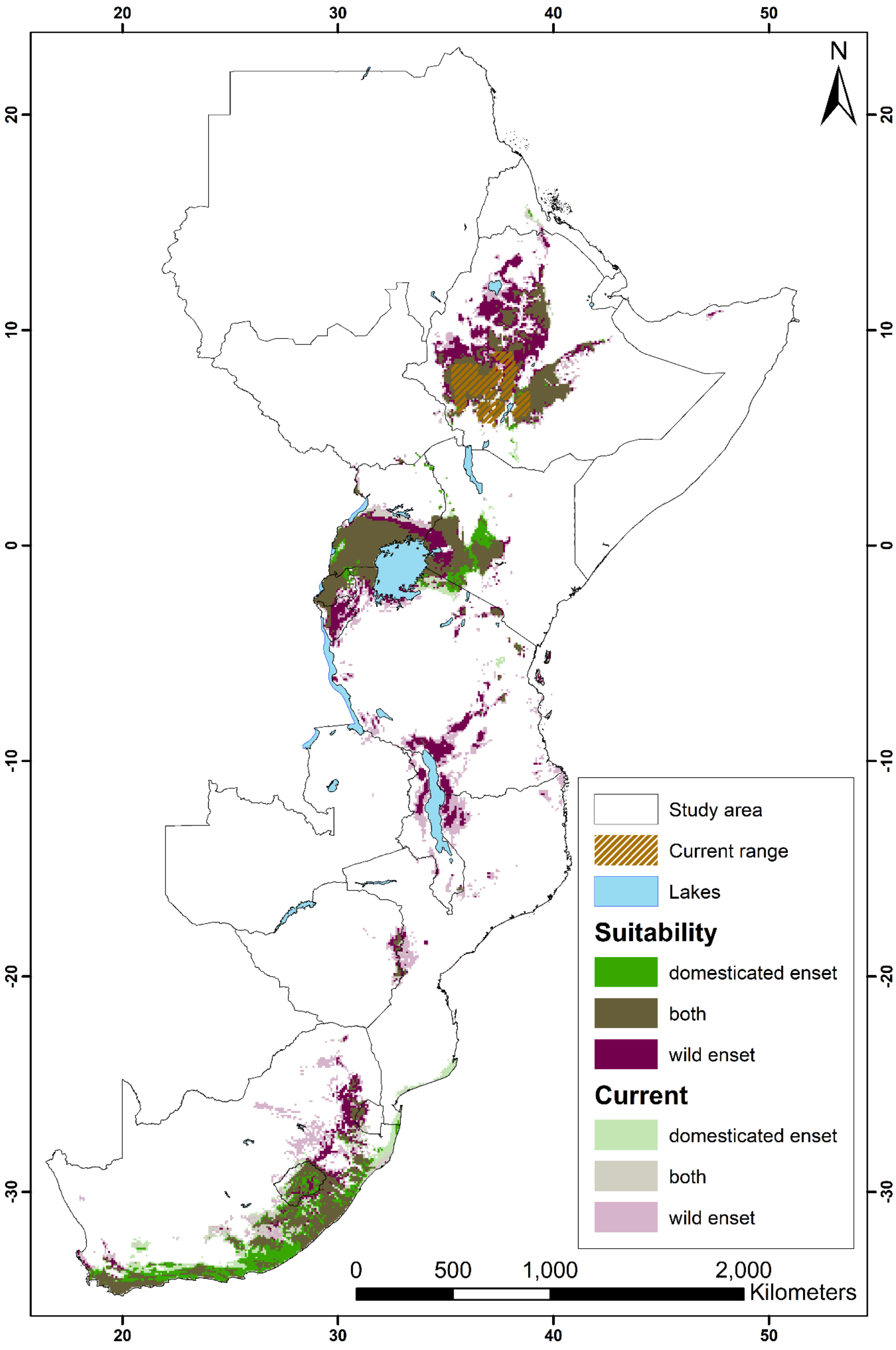

**Figure S4.** Suitability range change for wild and domesticated enset for 2050 under RCP4.5.

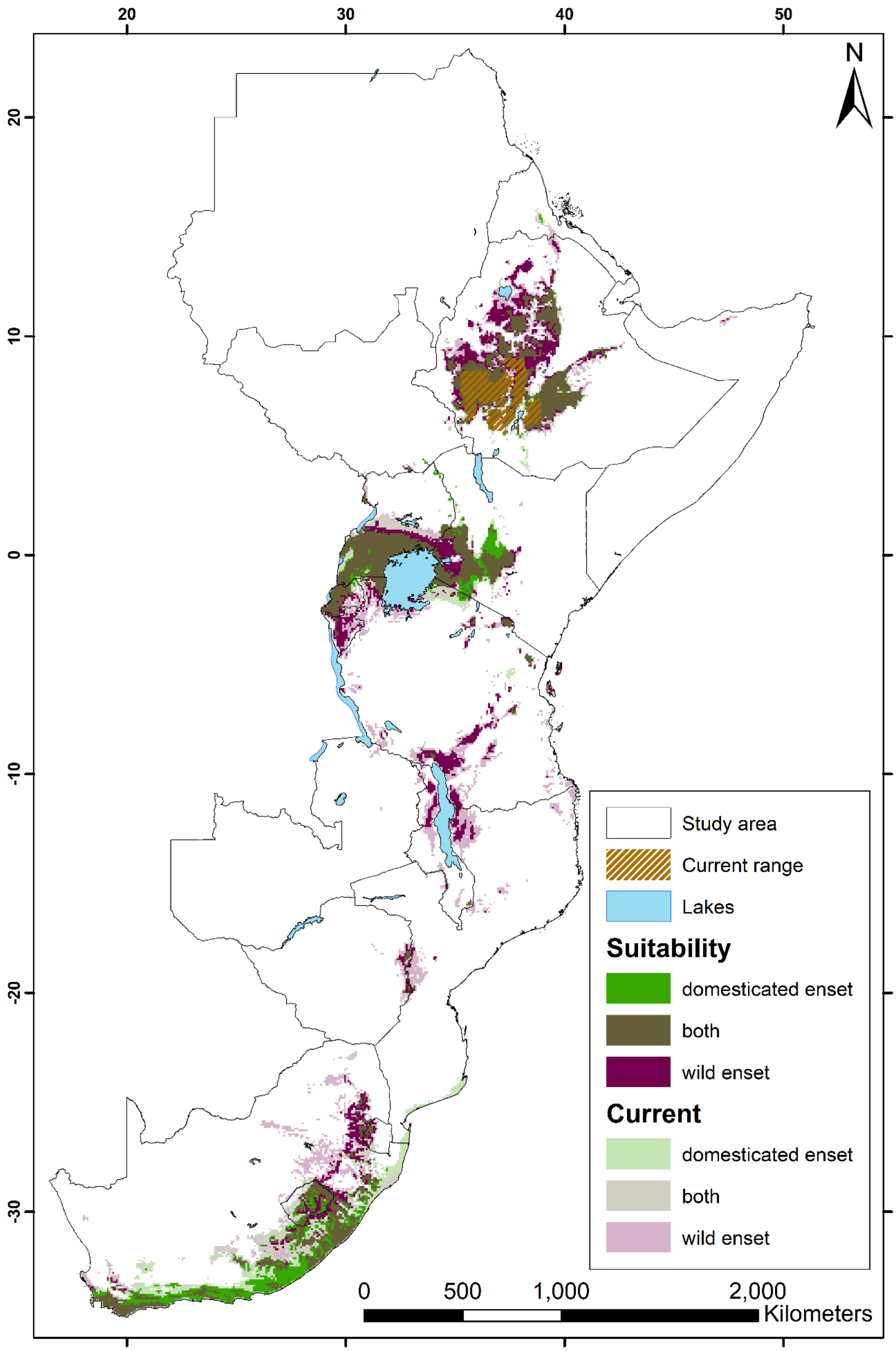

**Figure S5.** Suitability range change for wild and domesticated enset for 2050 under RCP8.5

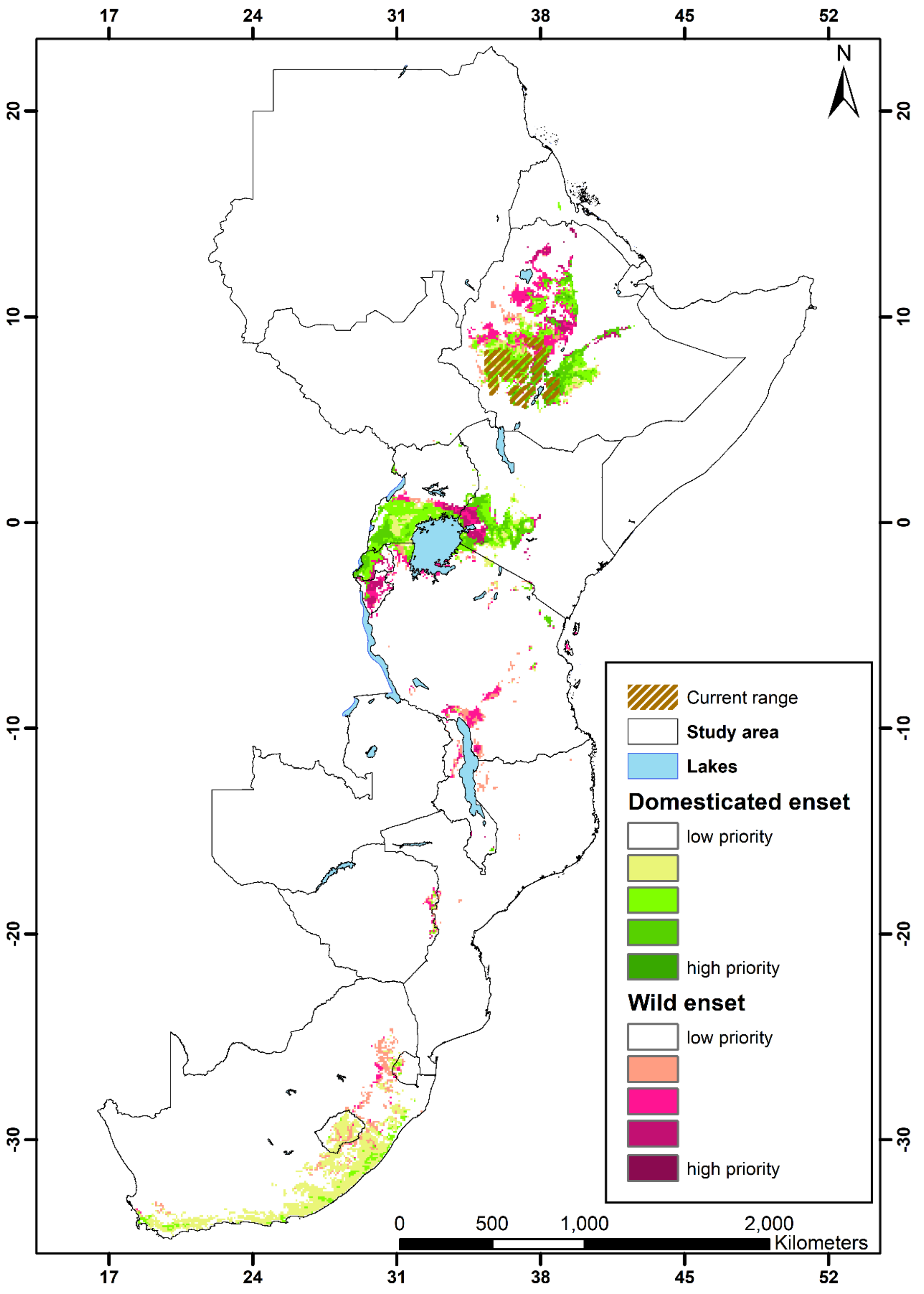

**Figure S6. Priority areas for enset cultivation in east Africa based on demographic and socioeconomic metrics** and enset suitability projections to 2070 under RCP 4.5 scenario. Both domesticated enset and the integration of crop wild relative diversity are considered

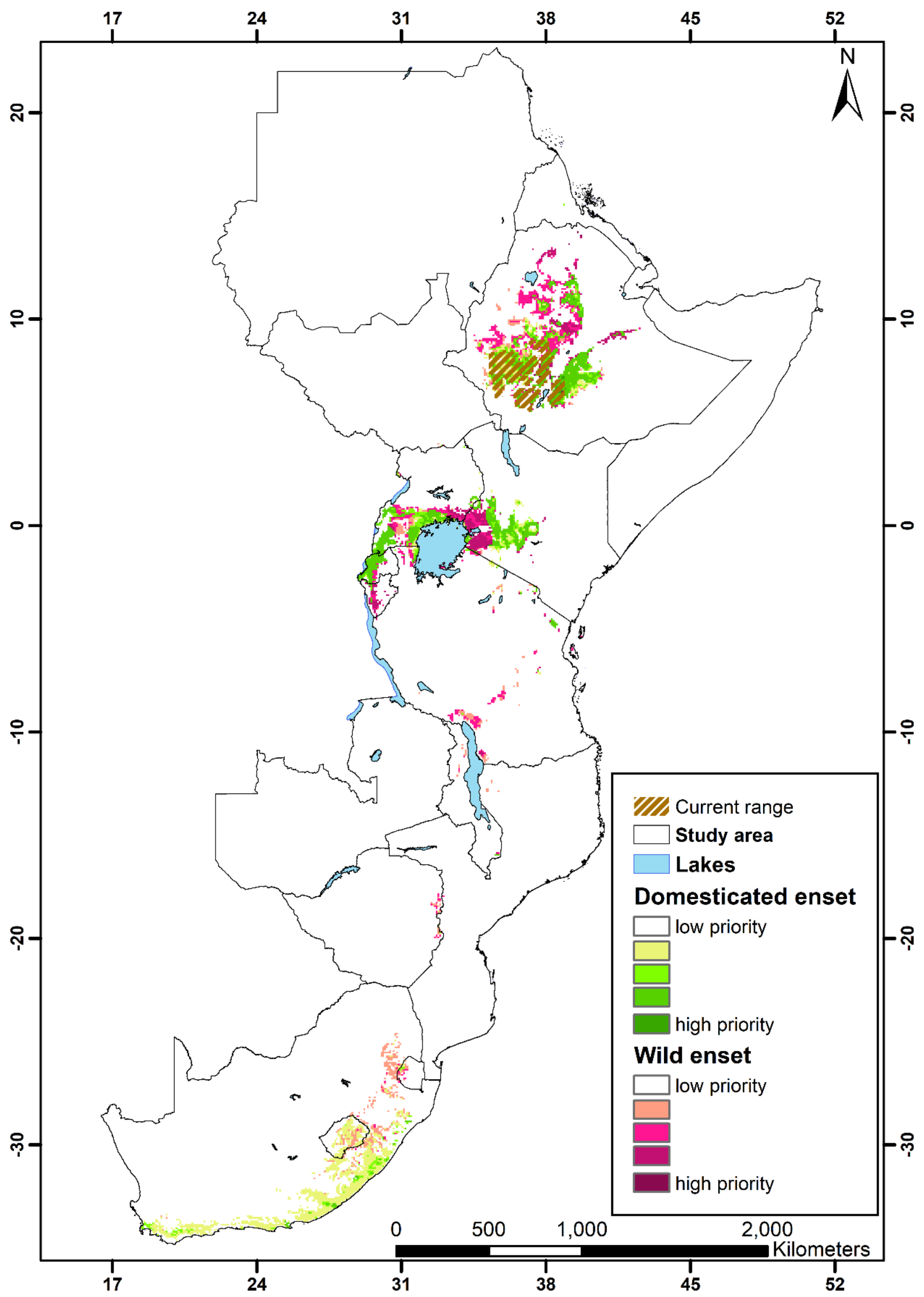

**Figure S7. Priority areas for enset cultivation in east Africa based on demographic and socioeconomic metrics** and enset suitability projections to 2070 under RCP 8.5 scenario. Both domesticated enset and the integration of crop wild relative diversity are considered.
